# Aberrant splicing in B-cell acute lymphoblastic leukemia

**DOI:** 10.1101/225136

**Authors:** Kathryn L. Black, Ammar S. Naqvi, Katharina E. Hayer, Scarlett Y. Yang, Elisabeth Gillespie, Asen Bagashev, Vinodh Pillai, Sarah K. Tasian, Matthew R. Gazzara, Martin Carroll, Deanne Taylor, Kristen W. Lynch, Yoseph Barash, Andrei Thomas-Tikhonenko

## Abstract

Aberrant splicing is a hallmark of leukemias with mutations in splicing factor (SF)-encoding genes. Here we investigated its prevalence in pediatric B-cell acute lymphoblastic leukemias (B-ALL), where SFs are not mutated. By comparing them to normal pro-B cells, we found thousands of aberrant local splice variations (LSVs) per sample, with 279 LSVs in 241 genes present in every comparison. These genes were enriched in RNA processing pathways and encoded ~100 SFs, e.g. hnRNPA1. hnRNPA1 3’UTR was pervasively misspliced, yielding the transcript subject to nonsense-mediated decay. Thus, we knocked it down in B-lymphoblastoid cells, identified 213 hnRNPA1-dependent splicing events, and defined the hnRNPA1 splicing signature in pediatric leukemias. One of its elements was DICER1, a known tumor suppressor gene; its LSVs were consistent with reduced translation of DICER1 mRNA. Additionally, we searched for LSVs in other leukemia and lymphoma drivers and discovered 81 LSVs in 41 genes. 77 LSVs were confirmed using two large independent B-ALL RNA-seq datasets. In fact, the twenty most common B-ALL drivers showed higher prevalence of aberrant splicing than of somatic mutations. Thus, post-transcriptional deregulation of SF can drive widespread changes in B-ALL splicing and likely contribute to disease pathogenesis.

## INTRODUCTION

Despite advances in the treatment of pediatric B-ALL, children with relapsed or refractory disease account for a substantial number of childhood cancer-related deaths. Adults with B-ALL experience even higher relapse rates and long-term event-free survival of less than 50% (Roberts and Mullighan, 2015). Recently, significant gains in the treatment of B-ALL have been achieved through the use of immunotherapies directed against CD19, a protein expressed on the surface of most B-cell neoplasms (Scheuermann and Racila, 1995; Sikaria et al., 2016). These gains culminated in the recent FDA approval of tisagenlecleucel and axicabtagene ciloleucel, CD19-redirected chimeric antigen receptor (CAR) T-cell immunotherapies, for patients with refractory/relapsed B-cell malignancies. However, relapses occur in 10-20% of patients with B-ALL treated with CD19 CAR T cells, often due to epitope loss and/or B-cell de-differentiation into other lineages (Gardner et al., 2016; Jacoby et al., 2016; Maude et al., 2014; Topp et al., 2015). Other targets for immunotherapy include CD20 and CD22 (Fry et al., 2017; Haso et al., 2013; Maino et al., 2016; Raetz et al., 2008). However, neither antigen is uniformly expressed in B-ALL, and factors accounting for this mosaicism are poorly understood (Sikaria et al., 2016).

We previously reported a new mechanism of pediatric B-ALL resistance to CD19-directed immunotherapy. We discovered that in some cases, resistance to CD19 CAR T cells was generated through alternative splicing of CD19 transcripts. This post-transcriptional event was mediated by a specific splicing factor (SF) SRSF3 and generated a CD19 protein isoform invisible to the immunotherapeutic agent via skipping of exon 2 ((Sotillo et al., 2015), reviewed by (Alderton, 2015; Behjati, 2015)).

Our discovery of a resistance mechanism based on alternative splicing prompted us to investigate the extent of this phenomenon in additional B-ALL cases. While driver mutations in splicing factors such as SRSF2, SF3B1, and U2AF1 have recently been discovered in myelodysplastic syndrome/acute myelogenous leukemia (Graubert et al., 2012; Papaemmanuil et al., 2011; Yoshida et al., 2011) and chronic lymphocytic leukemia (Quesada et al., 2012; Wang et al., 2011), SF mutations have not been reported in B-ALL. Nevertheless, our prior work suggested the possibility that SRSF3 (and by inference other SFs) could be deregulated in B-ALL (Sotillo et al., 2015), bringing about wide-spread splicing aberration.

This model would be particularly attractive because B-ALL is a chromosome translocation-driven disease where the prevalence of somatic mutations and copy number variations is relatively low. For example, the commonly mutated *IKZF1* gene (which encodes the Ikaros transcription factor) is affected by missense mutations in just ~20% of B-ALL cases. Similarly, mutations in the key tumor suppressor gene (TSG) TP53 are found in only ~7% of B-ALLs (per COSMIC database) (Forbes et al., 2015; Futreal et al., 2004). In addition, both genes are robustly transcribed across individual B-ALLs and thus are not epigenetically silenced. This raises the possibility that they and other TSGs are dysregulated by post-transcriptional events, such as alternative splicing.

## MATERIALS AND METHODS

### Bone Marrow Fractionation

Isolated mononuclear cells and whole bone marrow aspirates were obtained, respectively, from the University of Pennsylvania Stem Cell and Xenograft Core facility and CHOP Hematopathology Laboratory. For pediatric bone marrow samples, mononuclear cells were isolated by spinning over Ficoll gradient, as described earlier (Tasian et al., 2012). Residual red blood cells were lysed with Ammonium Chloride Lysis buffer with gentle rocking at room temperature for 10 min. Cells were pelleted by spinning at 250 x g for 10 min at 4° C and washed once with cold PBS/2%FBS. Cells were resuspended in 1mL PBS/2%FBS and incubated with 500uL FC Block on ice for 10 min. Cells were stained with 1mL CD34-PE, 500uL CD19-APC, and 500uL IgM-FITC for 30 min on ice. Cells were pelleted at 1300RPM for 6 min at 4° C and washed twice in cold PBS. Cells were resuspended in 3mL PBS/2%FBS containing 1uL/mL of 0.1mg/mL DAPI. Cells were sorted 4-ways using MoFlo ASTRIOS directly into RLT Lysis buffer (Qiagen) at a ratio of 1:3.

### Primary B-ALL Samples Acquisition

24 primary pediatric B-ALL samples were obtained from the CHOP Center for Childhood Cancer Research leukemia biorepository. Mononuclear cells from fresh bone marrow or peripheral blood specimens were purified via Ficoll gradient separation (Tasian et al., 2012) and cryopreserved for downstream experimental use.

### RNA-seq of bone marrow fractions, primary samples and cell lines

RNA was isolated using Qiagen RNeasy Mini Kit. RNA integrity and concentration were found using Eukaryote Total RNA Nano assay on BioAnalyzer (CHOP NAPCore). RNA-seq was performed on 10ng-1ug of total RNA according to GeneWiz protocol of PolyA selection, Illuminia Hi-seq, 2×150bp pair-end, 350M raw reads per lane.

### RNA-Seq alignment, quantification, and differential expression

Fastq files of RNA-seq obtained from GeneWiz were mapped using STAR aligner (Dobin et al., 2013). STAR was run with the option "alignSJoverhangMin 8". We generated STAR genome reference based on the hg19 build. Alignments were then quantified for each mRNA transcript using HTSeq with the Ensemble-based GFF file and with the option "-m intersection-strict". Normalization of the raw reads was performed using the trimmed mean of M-values (TMM). Principal component analysis (PCA) was done on normalized count values of the samples using a correlation matrix and a calculated score for each principal component. Differential expression of wild-type and knock-down or pro-B and B-ALL RNA-Seq datasets were assessed based on a model using the negative binomial distribution, a method employed by the R package DESeq2 (Love et al., 2014). Subsequent bar charts were generated using the R package ggplot2. Those differential genes that had a p-value of <0.05 were deemed as significantly up or down-regulated.

### Splicing analysis

In order to detect LSVs we used the MAJIQ (version 1.03) tool (Vaquero-Garcia et al., 2016). We ran MAJIQ on the Ensemble-based GFF annotations, disallowing *de novo* calls (Zerbino et al., 2017). We chose for further analysis LSVs that had at least a 20% change at a 95% confidence interval between two given conditions. Using in-house customized Perl and Ruby scripts, we filtered for events that corresponded to exon inclusion events only, forcing one event per LSV. For the genes containing differential LSVs in all 18 comparisons, we identified enriched gene ontologies (GO) using the gene functional classification tool DAVID (v. 6.7), reporting the top most significant hits based on p-value and false discovery rate. Heatmaps were generated for each ∆PSI value of each LSV that passed the 20% change threshold between Pro-B and B-ALL and additionally filtered for a frequency of n>=2 in our B-ALL samples. These were generated using the R package gplots. LSVs detected by MAJIQ were also tested using Leafcutter (version 0.2.7) (Li et al., 2018) for validation. We used LeafCutter for qualitative validation of LSV as it too, like MAJIQ, offers detection of complex and de-novo splice variants. However, LeafCutter’s model uses intron clusters that do not correspond directly to inclusion levels and is unable to model intron inclusion, hence assessment of exact inclusion levels or comparison to RT-PCR are generally not possible. In addition, we validated the splicing of hnRNPA1 3’UTR we performed RT-qPCR using a forward primer spanning exons 9-10 and reverse primers in exon 11. We normalized to actin and PHL156 as it most closely resembled Pro-B cells in its splicing pattern.

### Spearman’s Correlations

Correlations and their significance were computed using the nonparametric Spearman’s rank-order correlation implemented in R function *cor.test()*. Correlation analysis was performed between 1) RQ values of hnRNPA1 exon 11 inclusion as determined by RT-qPCR and PSI values of hnRNPA1 exon 11 inclusion as determined by MAJIQ, 2) RQ values of hnRNPA1 constitutive exons 2-3 and RQ values of hnRNPA1 exon 11 inclusion as determined by RT-qPCR (normalized to actin and PHL156), and 3) Normalized counts of hnRNPA1 constitutive exon 9. And normalized counts of hnRNPA1 exon 11 as determined by RNA-seq.

### Nonsense Mediated Decay Inhibition

To inhibit NMD we treated 2mil Nalm6 B-ALL lines in duplicate with either DMSO or 30ug/mL Cyclohexamide for 6 hours followed by RNA extraction. To confirm inhibition of NMD we performed gel-based PCR with primers spanning SRSF3 poison exon 4 to detect accumulation of a canonical NMD target. To determine if either hnRNPA1 3’UTR was an NMD substrate we performed RT-qPCR with forward primer spanning exons 9-10 and reverse primers in exon 11 or spanning exons 12-13. Data was normalized to hnRNPA1 constitutive exons 2-3 and DMSO.

### siRNA knock-down of hnRNPA1

Biological duplicate experiments were performed on 2 million P493-6 B-lymphoblastoid cells 27 electroporated using Amaxa Program O-006 with either 133nM non-targeting siRNA (Dharmacon) or 300nM ON-TARGET Plus Human hnRNPA1 SMARTpool siRNA (Dharmacon). Cells were plated in 2mL warm tetracycline-free RPMI for 24 hr. RNA isolation and RNA-seq were performed as described above. Knockdown was validated through RT-qPCR and Western Blot with anti-hnRNPA1 antibody (Abcam#ab5832). Pathway analysis was performed as described above. Our motif detection pipeline utilized RBPMap (v. 1.1) with a medium stringency level (p-value<=0.005) and conservation filter (Paz et al., 2014). We performed this analysis on exons affected and +/-200nts into adjacent intronic regions. We then identified hnRNPA1 motifs (Ray et al., 2013) and calculated the average z-score and p-value for all significant hits.

### Datasets

Cancer gene symbols and annotations were downloaded from the COSMIC database (Forbes et al., 2015; Futreal et al., 2004). Known splice factors were annotated and obtained from published studies or ensemble-annotated databases. Pediatric B-ALL samples from the TARGET consortium (phs000218.v19.p7) were accessed via NCBI dbGaP (the Database of Genotypes and Phenotypes) Authorized Access system. Pediatric B-ALL samples from the St Jude Children’s Research Hospital (EGAD00001002704 and EGAD00001002692) were accessed by permission from the Computational Biology Committee though The European Bioinformatics Institute (EMBL-EBI).

## RESULTS

### RNA-seq analysis of bone marrow-derived human B-cells

To determine if patterns of splicing dysregulation occur in B-ALL, we first generated normal B-lymphocyte datasets corresponding to potential cells of origin. To this end, we obtained from the University of Pennsylvania Stem Cell and Xenograft Core facility two healthy adult bone marrow biopsies and from the Children’s Hospital of Philadelphia (CHOP) Hematopathology Laboratory two bone marrow biopsies from children without leukemia. We enriched for mononuclear cells using Ficoll gradient separation, stained cells for combinations of stage-specific surface markers, and sorted B-cell subsets using flow cytometry. Specifically, we fractionated bone marrow progenitors into early progenitors (CD34^+^/CD19^-^/IgM^-^), pro-B (CD34^+^/CD19^+^/IgM^-^), pre-B (CD34^-^/CD19^+^/IgM^-^), and immature B (CD34^-^/CD19^+^/IgM^+^) populations (Fig. 1a top and Supp. Fig. 1a). We then extracted RNA, performed RNA-seq, and quantified transcript levels. Concordant with flow cytometric profiles, CD19 mRNA levels increased throughout differentiation stages, CD34 mRNA was confined to early progenitors and pro-B fractions, and IgM transcript was expressed only in the immature fraction (Fig. 1a, bottom). Furthermore, we performed principal component analysis (PCA) on all expression datasets and found tight clustering of the four fractions from different donors, suggesting that at the level of mRNA expression B-cell differentiation supersedes individual variations (Fig. 1b). To ensure that the clustering of fractions was not driven solely by CD19 and CD34 expression, we repeated the PCA, removing CD19 and CD34 expression contributions and observe very similar clustering of fractions (Supp. Fig 1b).

**Fig. 1.**
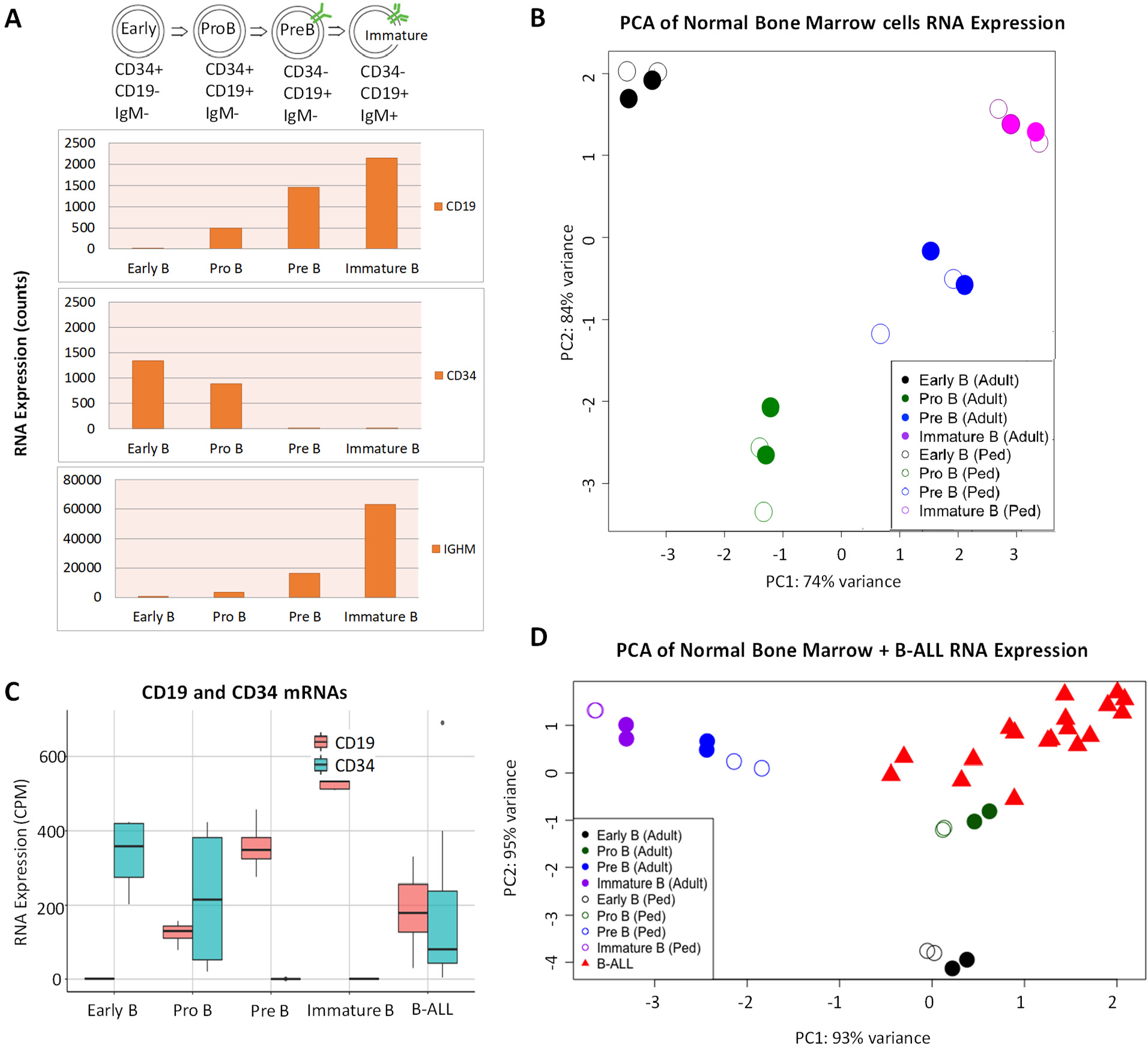
RNA-seq analysis of bone marrow-derived B-cell progenitors and pediatric B-ALL. a. Top: Lymphocytes were isolated from normal whole bone marrow aspirates and fractionated into CD34+/CD19-/IgM-, CD34+/CD19+/IgM-, CD34-/CD19+/ IgM-, and CD34-/CD19+/IgM+ populations. Bottom: Validation of surface marker expression by RNA-Seq in early progenitors, pro-B, pre-B, and immature B cells fractions. b. Principal component analysis (PCA) of RNA expression data from bone marrow fractions obtained from 2 adult (solid circles) and 2 pediatric (open circles) donors. Cumulative variances are shown for each PC. c. Quantification of CD19 and CD34 expression in bone marrow fractions and B-ALL samples. d. PCA on RNA expression data from 18 B-ALL and 4 bone marrow samples.

### RNA-Seq analysis of primary B-ALL samples

We obtained 24 primary pediatric B-ALL samples from the CHOP Center for Childhood Cancer Research leukemia biorepository. Mononuclear cells from fresh bone marrow or peripheral blood specimens were purified via Ficoll gradient separation and cryopreserved for downstream experimental use. We extracted total RNA and analyzed sample integrity. Based on the presence of intact RNA (evident by 28S and 18S bands on gels), an RNA integrity score >5.2, and RNA concentration of >40ng/uL, we successfully extracted high quality RNA for sequencing from 18 out of 24 frozen samples (Supp. Fig. 1c, red asterisks indicate samples that did not pass QC and were not sequenced). These samples were comprised of several different phenotypic and genetic B-ALL subtypes (Mullighan et al., 2007) (Supplementary Table 1 and Supp. Fig. 1d). We performed RNA-seq of these leukemia samples and first compared them to adult and pediatric normal bone marrow fractions with respect to CD19 and CD34 expression. We found that B-ALL samples closely resembled the pro-B fractions in that CD19 and CD34 transcripts were readily detectable in the low-to-medium range (Fig. 1c). To extend this analysis to the entire transcriptome, we then performed PCA on B-ALL samples and four sets of normal bone marrow fractions (two pediatric and two adult). Once again, the 18 B-ALL samples clustered most closely with the pro-B fractions (Fig. 1d). This similarity was reflected in shortest Euclidian distances between PC1 coordinates for the centroids of the B-ALL and pro-B cell samples (Supp. Fig. 1e). Therefore, we chose the pro-B fractions as cell-of-origin controls for B-ALL splicing analysis.

**Table 1.** Clinical and genetic characteristics of childhood B-ALL samples. USI = unique specimen identifier, D = diagnosis, R = relapse, F= female, M = male, SR = standard risk, HR = high risk.

| USI | Disease status* | Age at Dx, yrs | NCI Risk Status<sup>#</sup> | Sex | Race | Ethnicity | Major cytogenetic and molecular lesions |
| --- | --- | --- | --- | --- | --- | --- | --- |
| 11 | R | 7.5 | SR | F | Black or African American | Not Hispanic or Latino | high hyperdiploidy with +4 +10 |
| 50 | D | 11.7 | HR | M | White | Not Hispanic or Latino | 46,XY, IKZF1 deletion |
| 54 | D | 3.9 | SR | M | Other | Hispanic or Latino | ETV6-RUNX1, PAX5 deletion |
| 102 | D | 6.6 | SR | F | American Indian or Alaska Native | Not Hispanic or Latino | ETV6-RUNX1 |
| 84 | D | 16.1 | HR | F | Black or African American | Not Hispanic or Latino | high hyperdiploidy |
| 84 | R |  |  |  |  |  | high hyperdiploidy, CREBBP deletion |
| 105 | D | 2.4 | SR | F | White | Not Hispanic or Latino | high hyperdiploidy with +4 +10 |
| 145 | D | 7.7 | HR | M | Other | Hispanic or Latino | EBF1-PDGFRB, IKZF1 deletion |
| 149 | D | 15.0 | HR | F | Other | Not Hispanic or Latino | 45,XX , CDKN2A/B deletion |
| 151 | D | 8.5 | SR | F | White | Not Hispanic or Latino | CDKN2A deletion, IKZF1 deletion, ERG deletion |
| 156 | D | 6.3 | HR | F | Other | Not Hispanic or Latino | high hyperdiploidy, iAMP21 |
| 158 | D | 3.5 | SR | F | White | Not Hispanic or Latino | ETV6-RUNX1 |
| 161 | D | 3.9 | HR | M | White | Not Hispanic or Latino | 47,XY +22, iAMP21, IKZF1 deletion |
| 163 | D | 5.6 | SR | M | White | Not Hispanic or Latino | ETV6-RUNX1, PAX5 deletion |
| 222 | D | 16.0 | HR | M | Black or African American | Not Hispanic or Latino | 46,XY, IKZF1 deletion, |
| 540 | D | 1.9 | HR | F | White | Not Hispanic or Latino | ETV6-RUNX1 |
| 1074 | D | 5.2 | SR | F | White | Not Hispanic or Latino | ETV6-RUNX1, CDKN2A/B deletion |
| 1200 | D | 5.6 | SR | M | White | Not Hispanic or Latino | ETV6-RUNX1, NRAS Q61H; germline +21 |

### Global patterns of aberrant splicing in pediatric B-ALL

To detect patterns of alternative splicing in B-ALL, we utilized the MAJIQ (Modeling Alternative Junction Inclusion Quantification) algorithm (Vaquero-Garcia et al., 2016). MAJIQ offers the ability to detect, quantify, and visualize complex splicing variations, including de-novo variations, while comparing favorably in reproducibility and false discovery rate to common alternatives (Norton et al., 2017; Vaquero-Garcia et al., 2016). Using MAJIQ, we performed 18 independent pairwise comparisons between the average of pediatric pro-B cell fractions and leukemia samples. To assess heterogeneity in splicing across samples, we measured the number of differential LSVs (minimum of 20% change in splicing, 95% CI) in each sample and compared their identities. We observed that each B-ALL specimen had >3000 LSVs but most LSVs were either unique (first bar in Fig. 2a) or shared by only a small number of B-ALL patient samples (subsequent bars in Fig. 2a). However, we also found a total of 279 aberrant LSVs in 241 genes that were detected in all of our 18 pairwise comparisons (last bar in Fig. 2a). Provocatively, when this 241-gene set was analyzed using DAVID (Database for Annotation, Visualization, and Integrated Discovery) (Dennis et al., 2003), the top gene ontology (GO) categories (Blake et al., 2013) were related to RNA splicing (Fig. 1a, inset). When we investigated individual SF transcripts, we found that of the 167 well-characterized SF genes (Supplementary Table 2, from (Sebestyen et al., 2016)), 101 were alternatively spliced in B-ALL compared to pro-B cells (Fig 2b, bottom left, proB- vs B-ALL bar chart). Moreover, these changes were highly specific to the malignant phenotype and not to states of normal B-cell differentiation (Fig 2b, upper and middle panels). For example, only 6 SF transcripts were alternatively spliced during the early progenitors-to-proB transition and only 9 during the proB-to-preB transition (Fig 2b middle).

**Table 2.**
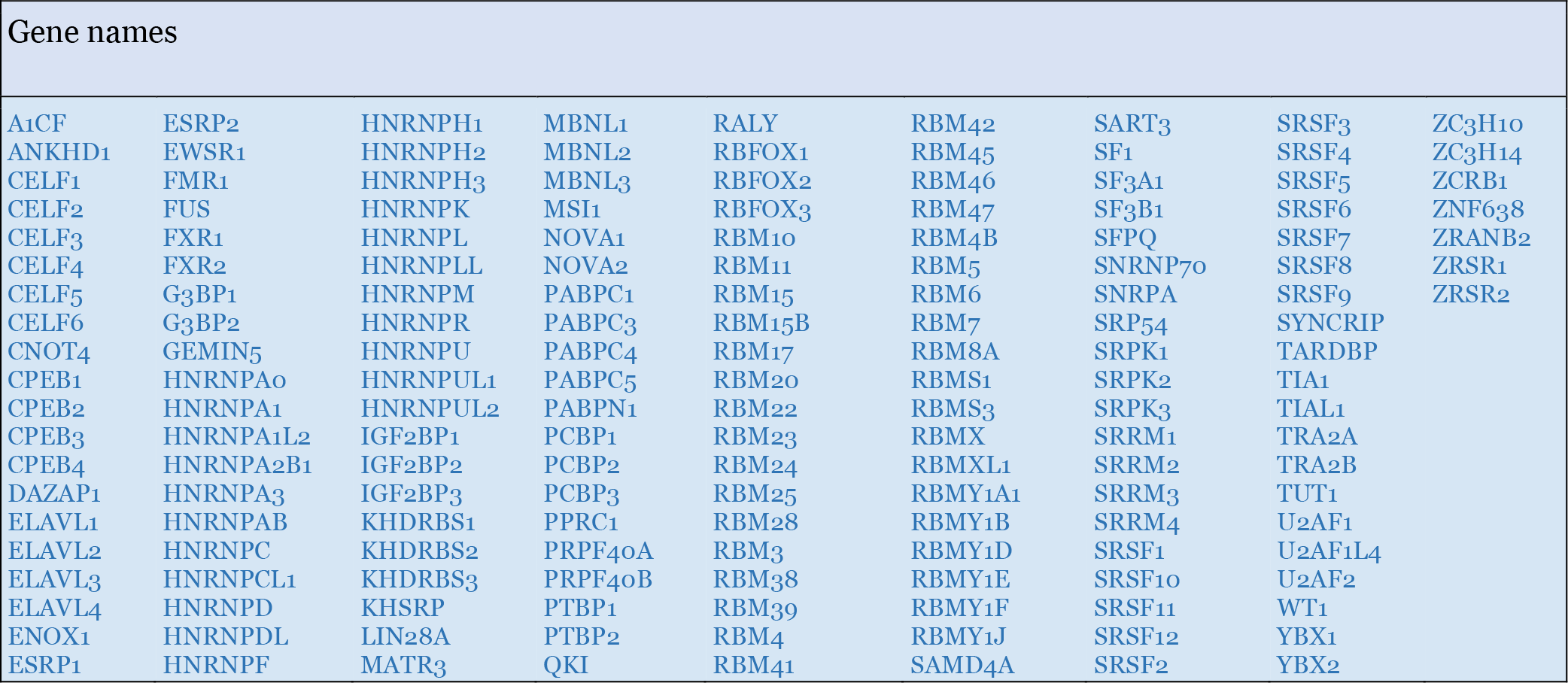
Well characterized splicing factors analyzed for aberrant LSVs.

**Fig. 2.**
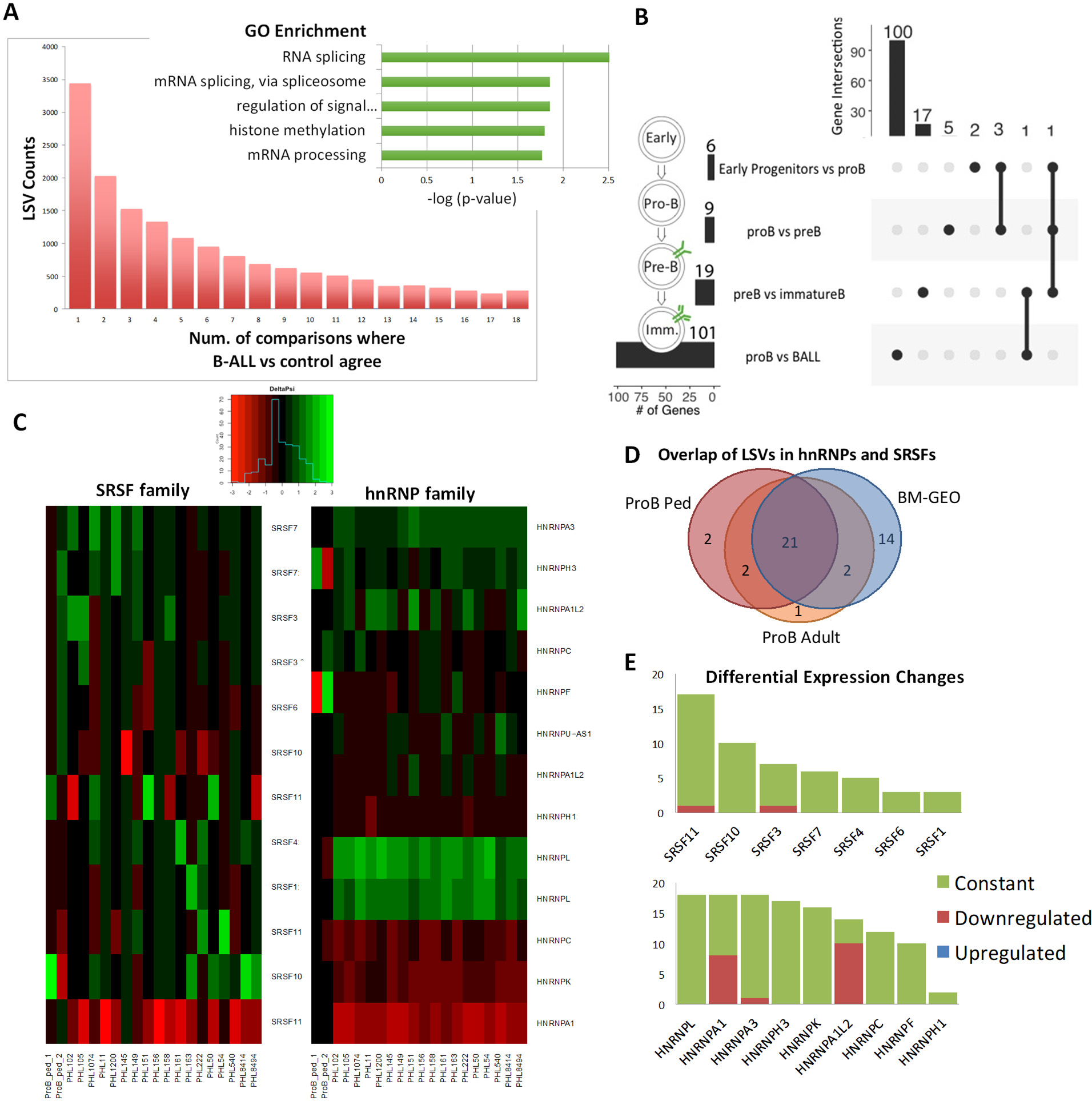
Alternative splicing in pediatric B-ALL. a. Enumeration of local splicing variations (LSVs). The vertical bar chart shows the numbers of LSVs detected by MAJIQ per B-ALL sample compared to normal pro-B cells. The horizontal bar chart (insert) showing enrichment for RNA splicing pathways among genes with most consistent LSVs. b. Overlap of LSVs in SF genes between B-ALL samples and bone marrow fractions. The left panel shows the different sets and number of genes included in them. The middle section visualizes the different combinations of intersection (including each set by itself). The upper panel shows how many genes of the total set are in each intersection. c. Heat maps showing changes in inclusion of exons corresponding to SRSF and hnRNP families members in 18 B-ALL samples compared to the pro-B cell fraction. Green color denotes increased inclusion, red color – increased skipping. d. Venn diagram showing overlap of LSVs from hnRNP and SRSF families when B-ALL samples are compared to in-house pediatric pro-B, in-house adult pro-B, and publicly available bone marrow data (GSM1695856 and GSM1695857), as indicated. e. Differential expression analysis of hnRNP and SRSF family members to decouple detection of LSVs with changes in RNA expression. Majority of samples showed constant expression of indicated genes compared to normal pro-B samples (green). Several samples showed downregulation of a few genes (red), notably hnRNPA1.

For each LSV, MAJIQ provides a ∆PSI (percent-spliced-in) value to indicate changes in isoform abundance. We observed that many members of the hnRNP and SRSF families, which play key roles in exon inclusion or skipping (Matera and Wang, 2014), exhibit profound changes in ∆PSI values (Fig 2c). These events included increases in the so-called ‘poison exons’ with in-frame stop codons in several SRSF proteins (Jumaa and Nielsen, 1997). Of note, some transcripts contain multiple aberrant LSVs. For example, SRSF11 has three distinct LSVs with different corresponding start/end genomic coordinates.

To increase confidence in the detection of splice factor related LSVs in B-ALL compared to normal pediatric pro-B samples we also compared B-ALL to normal adult pro-B cells and to publicly available RNA-seq data corresponding to the CD19-positive normal bone marrow cells (Casero et al., 2015). We then looked for overlap between each of these comparisons and detected 21 out of 25 LSVs consistently altered in B-ALL when compared to normal pediatric pro-B, adult pro-B or bone marrow samples (Fig 2d).

To decouple observed changes in splicing from changes in gene expression we performed differential expression analysis on genes with frequently detected LSVs. We found that most genes containing LSVs did not have associated changes in expression, with the notable exception of hnRNPA1, which was observed to be downregulated (two-fold difference) in approximately half of the B-ALL samples (Fig 2e). This and the pervasive nature of hnRNPA1 splicing alterations (Fig 2d, right) prompted us to investigate its regulation further.

### Dysregulated splicing of hnRNPA1 in B-ALL

According to the Ensembl database (Zerbino et al., 2017), there is evidence for hnRNPA1 transcripts with alternatively spliced 3’ UTRs, notably the canonical long proximal exon 11 and two shorter, distal exons (exons 12/13, ENST00000547566) (Fig. 3a, top). We observed that the exon 11-containing transcript predominated in normal pro-B cells (Fig 3b, red), but in a typical B-ALL sample (representative sample PHL50) the predominant event was the skipping of exon 11 to exons 12 and 13 (Fig. 3b, blue). To ensure validity of this event, we also ran the LeafCutter splicing algorithm to detect splice events present in the averaged RNA-seq datasets from 18B-ALL samples and 4 pro-B samples. Confirming our MAJIQ analysis we were able to detect alternative splicing of hnRNPA1 3’UTR with this independent analysis (Supp. Fig. 2a). While exon 11 percent-spliced-in (PSI) values varied across leukemia samples, most leukemias had increased skipping of exon 11 compared to pro-B samples (Fig. 3c, blue). We also observed intron 10 inclusion in all pro-B and B-ALL samples, although this event was not significantly different between normal and malignant samples (Fig 3c, grey stacks).

**Fig. 3.**
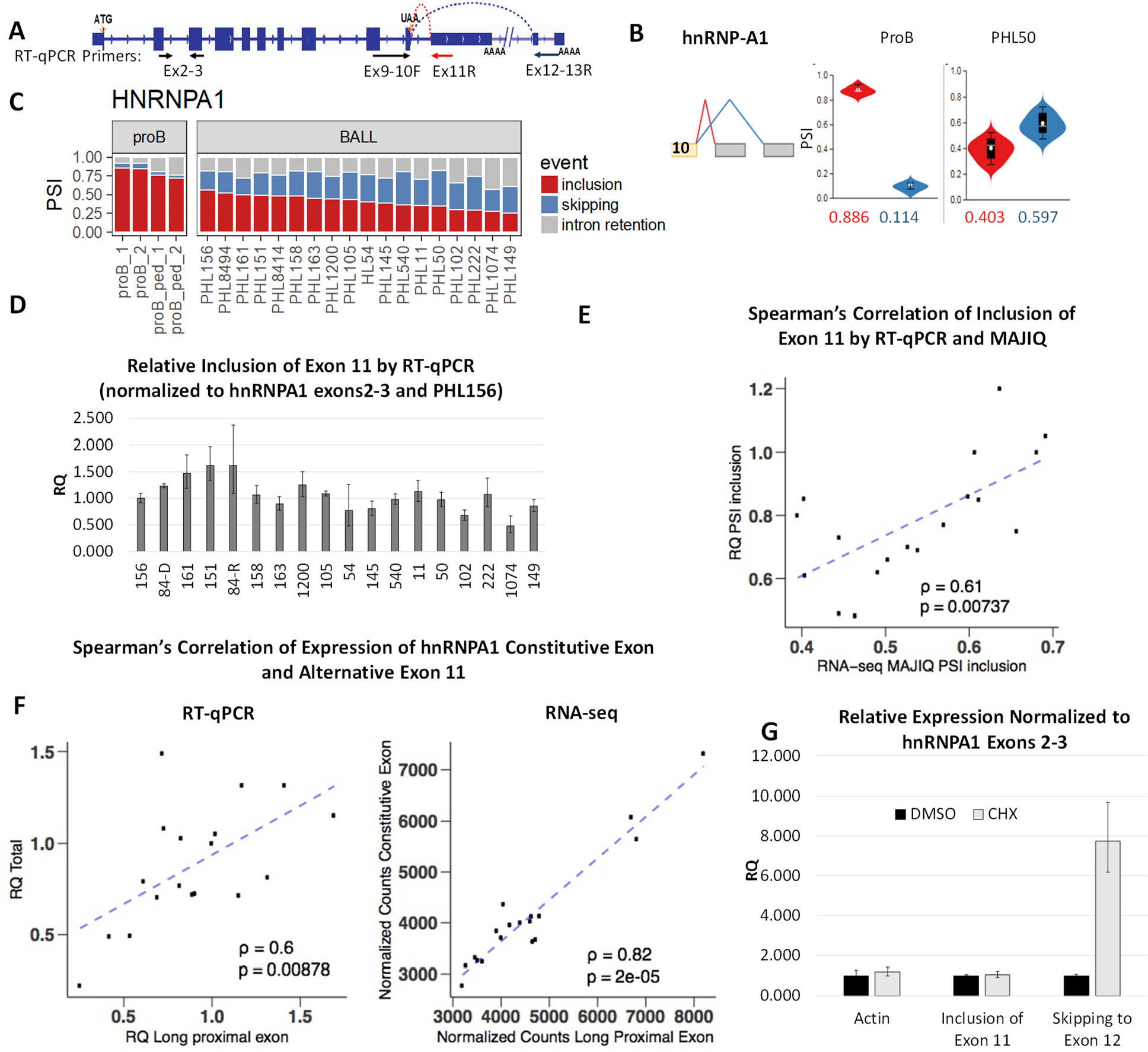
Alternative splicing of hnRNPA1 in pediatric B-ALL. a. Exon-intron structure of the HNRNPA1 transcript and location of RT-qPCR primers. b. Violin plots depicting alternative splicing of the hnRNPA1 transcript. c. Stack plot of PSI values of exon 11 corresponding to pro-B cell fractions and 18 B-ALL samples. d. Analysis of exon 11 inclusion in B-ALL samples by RT-qPCR. e. Spearman’s correlation of RT-PCR vs. MAJIQ (rho=0.06, p=0.00737). f. Spearman’s correlations of expression of an hnRNPA1 constitutive exon and alternative exon 11 measured by RT-qPCR (left, rho=0.6, p=0.00878) or RNA-Seq (right, rho=0.82, p=2E-05). g. Relative expression of actin and alternative hnRNPA1 3’UTR exons 11 or 12 in the presence of DMSO (black) or NMD-inhibitor cyclohexamide (CHX, grey).

We then validated and quantitated exon 11 inclusion by RT-qPCR using a forward primer spanning exons 9-10 and a reverse primer in exon 11 (Fig 3a,d). Using Spearman’s *rho* statistic we observed a strong positive association between MAJIQ predictions and RT-qPCR validations (0.61, p-value=0.00737) (Fig. 3e). We next applied the same correlation analysis to hnRNPA1 mRNA expression levels versus exon 11 inclusion and found a positive correlation between the two measurements when measured by RT-qPCR (0.6, p-value=0.00878) or RNA-seq (0.82, p-value=0.00002) (Fig. 3f). This suggested that preferential splicing from exon 10 to the distal 3’ UTR exons 12/13 (skipping exon 11) may decrease hnRNPA1 RNA steady state levels. Based on the exon junction present between exons 12 and 13 present downstream from the stop codon we speculated this transcript might be a target of NMD. To test this, we treated Nalm6 B-ALL cells with translation inhibitor cyclohexamide to transiently inhibit NMD (Amrani et al., 2004; Carter et al., 1996; Huang et al., 2011). To show effective inhibition of NMD, we performed gel-based PCR for primers spanning SRSF3 exon 4, a canonical target of NMD (Supp Fig 2b). We then performed RT-qPCR for hnRNPA1 3’ UTR exons and observed accumulation of the transcript that skips to exon 12/13, but not actin or the transcript including exon 11 (Fig 3g). This indicated that transcripts that skip exon 11 to exons 12/13 may be indeed targeted by NMD.

To model this event and identify potentially affected transcripts, we knocked down hnRNPA1 with siRNA in the Epstein-Barr Virus (EBV) transformed human B-lymphoblastoid P493-6 cell line (Pajic et al., 2001) and performed RNA-seq analysis. Knockdown of hnRNPA1 was confirmed by RT-qPCR (Fig 4a) and western blot (Fig 4b). Differential expression analysis showed that levels of hnRNPA1 mRNA were robustly decreased by the hnRNPA1 siRNA compared to the non-targeting siRNA control (Supp. Fig. 2c). Using MAJIQ we identified 213 LSVs in 184 genes associated with knockdown of hnRNPA1 (minimum of 20% ∆PSI, 95% CI). Of these hnRNPA1-dependent LSVs, 74 (or more than 30%) were present in at least one B-ALL sample (Fig. 4c), with more than half of these 74 LSVs altered in 10 or more samples. DAVID analysis of the genes with overlapping LSVs revealed enrichment for macromolecular metabolic pathways (Fig 4d), supporting the well-established role of hnRNP proteins in RNA metabolism (Weighardt et al., 1996). We next searched for presence of the hnRNPA1 binding motif (Burd and Dreyfuss, 1994) in the exons and flanking intronic sequences (+/-200nt) involved in overlapping LSVs. We identified the UAGGG motif in 68 out of 74 LSVs in question. Among these genes characterized by overlapping LSVs, involvement in macromolecular metabolism, and presence of hnRNPA1 binding motif, was DICER1. Interestingly, one of the LSVs in the DICER1 gene was altered in B-ALL samples in the same manner as in the hnRNPA1 KD cells (Fig. 4f). We also confirmed the presence of this LSV using LeafCutter algorithm (Supp. Fig2d). This LSV maps to the 5’UTR of the DICER1 transcript (Supp. Fig. 2e), potentially diminishing its translation efficiency. This type of deregulation would be consistent with the loss-of-function mutation and copy number variations in DICER1 in several types of cancer including leukemias (Foulkes et al., 2014).

**Fig. 4.**
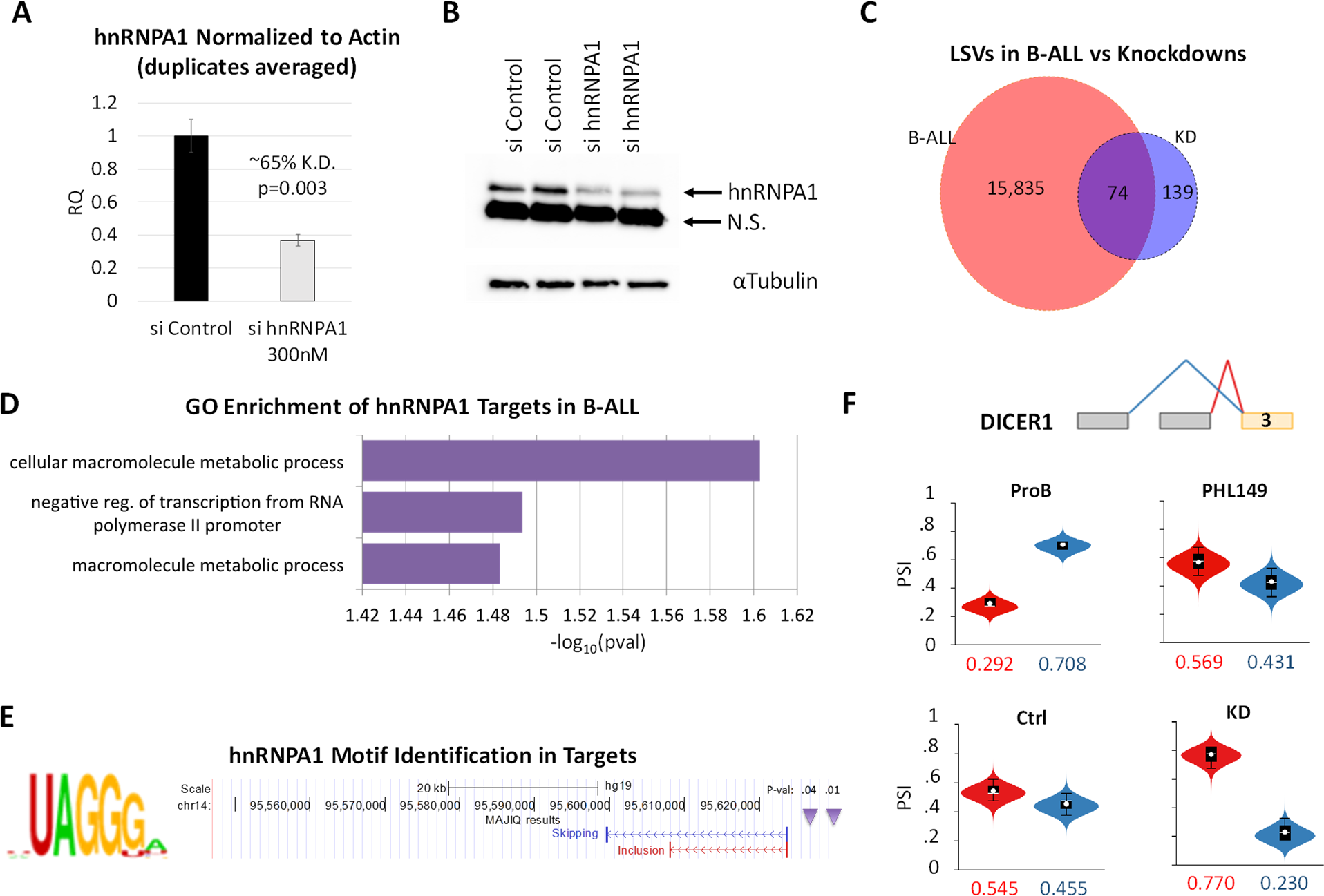
Alternative splicing of hnRNPA1 in pediatric B-ALL. a. Expression of hnRNPA1 RNA in P493-6 cells as determined by RT-qPCR (ex2-3 primers). Non-targeting control (black) and hnRNPA1-targeting (black) siRNA were used in duplicate experiments. b. hnRNPA1 protein expression determined by immunoblotting with antihnRNPA1 and anti-tubulin antibodies in cells from panel a. c. Overlap in LSVs detected in B-ALL versus pro-B and hnRNPA1 knockdown versus control cells. d. The horizontal bar chart showing enrichment for macromolecular metabolic pathways among genes with LSVs detected in both hnRNPA1 knockdown and B-ALL. e. hnRNPA1 Motif analysis of exons and adjacent intronic sequences (+/-200nt) involved in LSVs detected in hnRNPA1 knockdown and B-ALL samples. f. Violin plots depicting alternative splicing of DICER1 in Pro-B vs. PHL149 B-ALL (top) and hnRNPA1 control and knockdown P493-6 cells (bottom).

### Aberrant splicing of leukemia drivers in pediatric B-ALL

We next aimed to identify alternative splicing in other genes that contribute to leukemogenesis. We retrieved all genes (141) with known somatic mutations in hematologic malignancies from the COSMIC database (v.82) (Forbes et al., 2015; Futreal et al., 2004) (Supplementary Table 3) and searched for aberrant LSVs affecting these genes in B-ALL samples using the very stringent ΔPSI>50% cutoff for exon inclusion/skipping. We identified 81 aberrant LSVs in 41 genes that were present in at least two B-ALL samples. They accurately separated leukemia samples from the non-transformed pro-B cell counterparts following hierarchical clustering using Euclidian distances (Fig. 5a). Of note, these LSVs affected roughly a third of B-ALL driver genes. Some (in genes such as FBXW7) were present in almost all B-ALL samples, attesting to their potential significance in leukemia pathogenesis. Again, to decouple changes in splicing from changes in expression we performed differential expression analysis on genes containing LSVs in the COSMIC genes. Consistent with SF data analysis, the majority of genes in the majority of samples did not have changes in expression associated with changes in splicing (Supp. Fig. 3a).

**Table 3.**
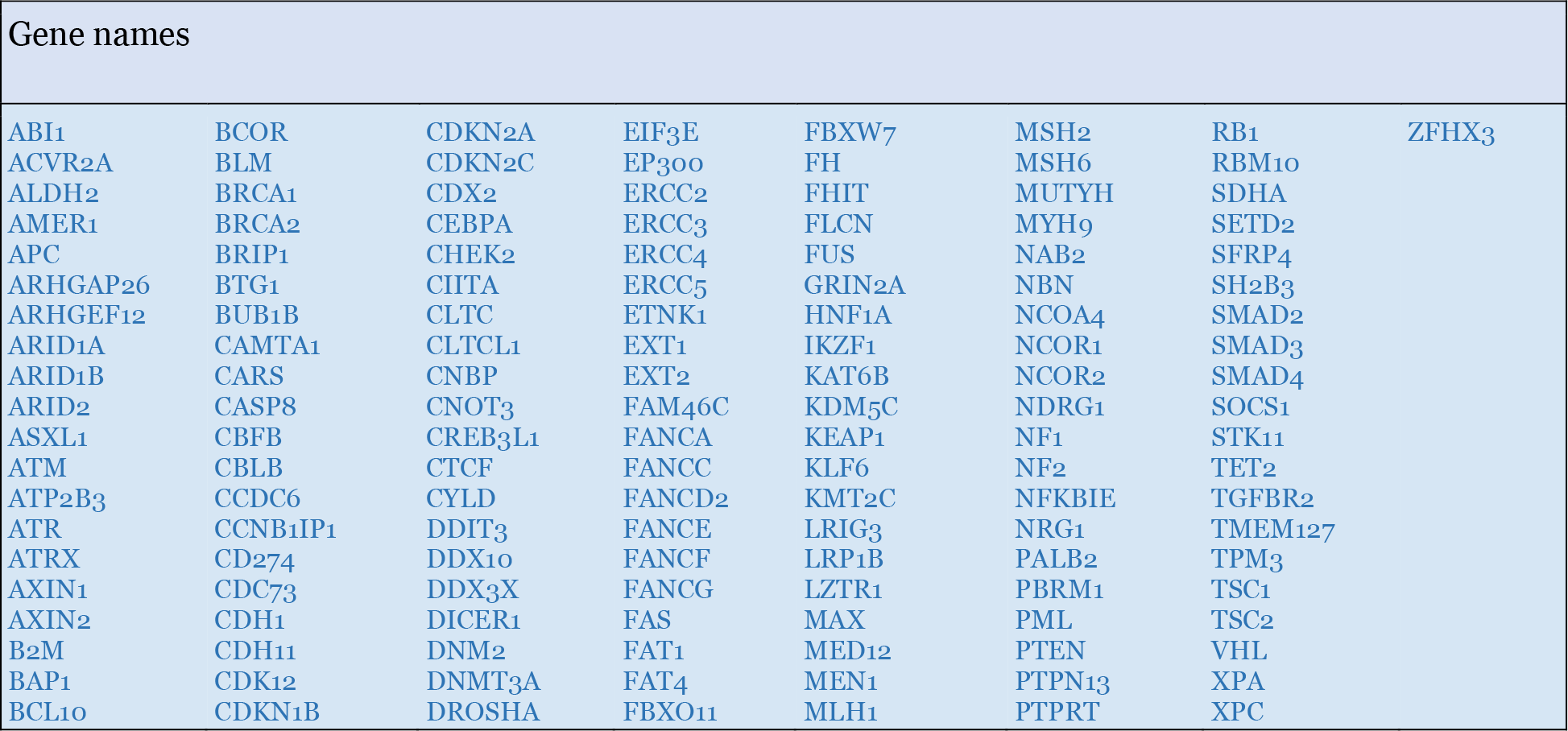
COSMIC genes mutated in hematologic malignancies and analyzed for aberrant LSVs.

**Fig. 5.**
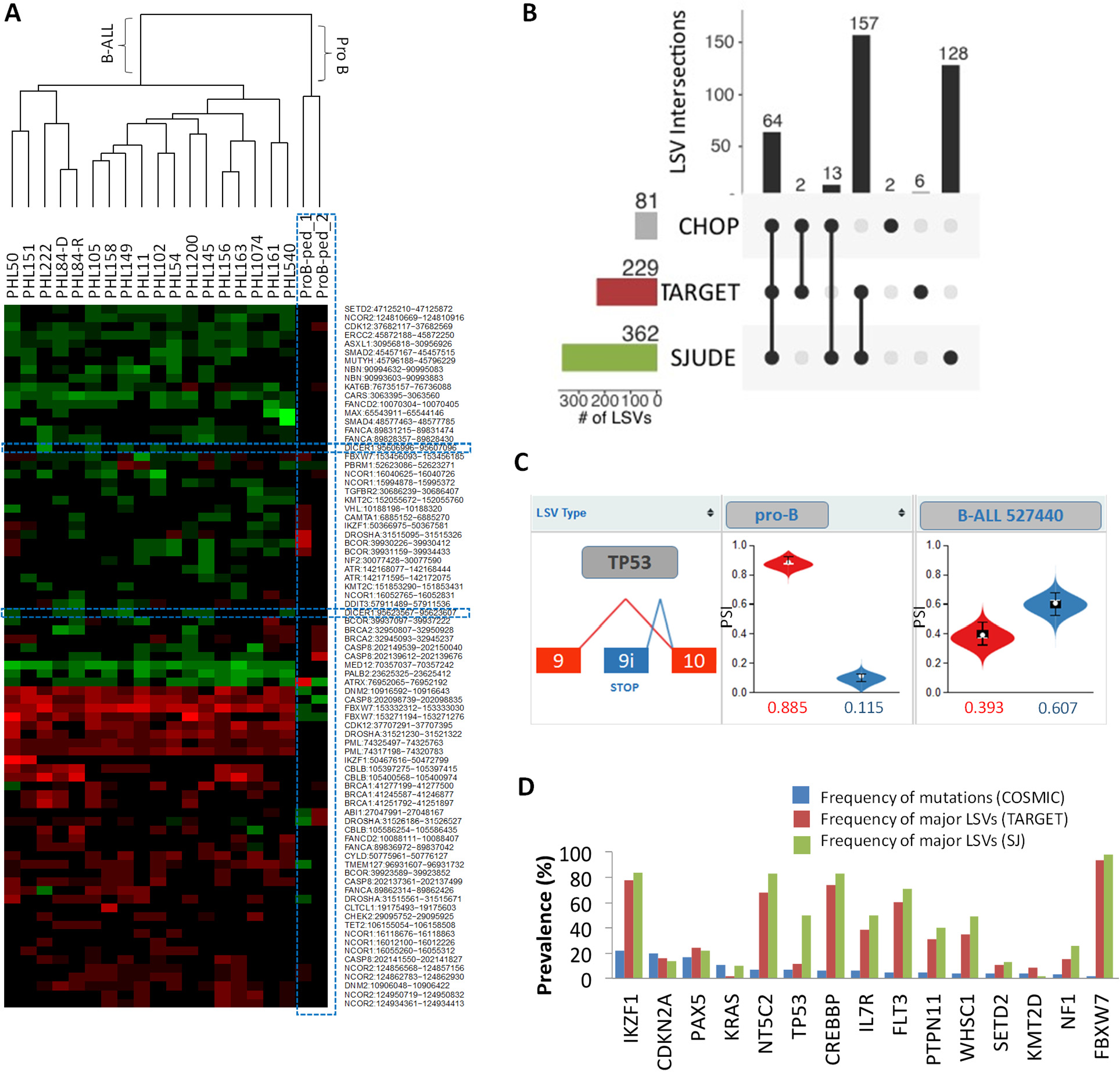
Alternative splicing of leukemia drivers in pediatric B-ALL. a. Heat maps showing changes in splicing in B-ALL of genes commonly mutated in hematologic malignancies. The dendrogram on top of the heatmap represents results of hierarchical clustering. The dotted blue line denotes the hnRNPA1-dependent LSV in DICER1 transcript. b. Overlap between the LSVs from panel (a) and those detected in TARGET and SJCRH B-ALL datasets. c. Violin plots depicting alternative splicing of TP53 in Pro-B and a representative TARGET B-ALL sample indicating increased inclusion of alternative exon 9i (blue) in B-ALL samples. d. Bar graph representing frequencies of mutations (COSMIC dataset) and major LSVs as defined by MAJIQ in the indicated datasets.

To confirm our findings in independent datasets, we searched for aberrant LSVs in the COSMIC genes in the TARGET (Hunger et al., 2013) and St Jude Children’s Research Hospital (SJCRH) (Gu et al., 2016) B-ALL datasets using pediatric pro-B cells for comparison. We found 229 LSVs in 80 genes in the TARGET data and 362 LSVs in 95 genes in the SJCRH data (Fig 5b). Importantly, 64 of 81 LSVs identified in CHOP datasets significantly overlapped with TARGET (Fisher’s Exact Test, p=7e-124) and SJCRH datasets (Fisher’s Exact Test, p=1e-139), 13 different LSVs are present in both CHOP and SJCRH datasets and another 2 LSVs are present in CHOP and TARGET datasets (Fig 5b). We then narrowed our analysis to LSVs in top 20 B-ALL tumor suppressors and oncogenes as defined in the COSMIC database. In both datasets, we discovered largely overlapping LSVs corresponding to 15 out of 20 genes, such as IL7R, FLT3, TP53, etc. For example, we detected increased inclusion of TP53 exon 9i in B-ALL compared to pro-B (Fig. 5c, red), which would promote the expression of the SRSF3-regulated p53-β isoform (Supp. Fig. 3b) involved in cell cycle and cell death (Bourdon et al., 2005; Fujita et al., 2009; Marcel et al., 2014; Tang et al., 2013). Of note, LSV frequencies were much greater than frequencies of somatic mutation (Fig. 5d), suggesting that in B-ALL these driver genes are preferentially affected by post-transcriptional mechanisms, such as alternative splicing.

## DISCUSSION

Personalized cancer diagnostics traditionally employ selected oncogene panels, which can identify mutations in specific genes known or suspected to be drivers in human malignancies. Hematologic malignancy sequencing panel typically include dominant oncogenes (e.g., FLT3 and IL7R), recessive tumor suppressors (e.g., TP53 and FBXW7), and haploinsufficient DNA/RNA caretakers (e.g., splicing factor SRSF2). Results from such genetic profiling of diagnostic cancer specimens can identify prognostic mutations and inform treatment selection for patients. Our data demonstrate that genetic deregulation occurs in B-ALL at the level of splicing in the absence of genetic mutations. For example, our analyses of B-ALL transcriptomes demonstrated that several SRSF genes controlling exon inclusion (Matera and Wang, 2014) show widespread variations in their own splicing patterns, some of which are known to decrease protein levels (Jumaa and Nielsen, 1997). Consequently, we observed aberrant splicing of some of their known target transcripts such as TP53 (Tang et al., 2013), which encodes the key tumor suppressor gene. Thus, our current data show that clinical genetic testing panels may be inadequate to identify all potential therapeutic vulnerabilities within B-ALL cells. Of note, the majority of aberrant LSVs were highly concordant across different datasets. This reproducibility validates our conclusions and alleviates the potential concern that sample preparation conditions (e.g., storage at ambient temperature) could be affecting RNA surveillance and thus impacting analysis of alternative splicing (Dvinge et al., 2014).

One of the most consistent changes in exon usage we observed was non-canonical selection of 3’ UTRs of hnRNPA1, a splice factor implicated in cancer progression (David et al., 2010; Michlewski and Caceres, 2010). While it is overexpressed in some cancers, in the B-ALL model, hnRNPA1 LSV correlated with a decrease in hnRNPA1 mRNA abundance. This is in agreement with our data that suggests this multi-exon 3’UTR would trigger non-sense mediated decay of the transcript. It is also possible that new 3’UTR sequences could create additional sites for targeting by microRNAs, which are known to play a key role in hnRNPA1 downregulation in chemotherapy-resistant ovarian cancer cells (Rodriguez-Aguayo et al., 2017). Interestingly, knockdown of hnRNPA1 in an Epstein-Barr Virus (EBV) transformed human B-cell lymphoblastoid cell line resulted in aberrant splicing of Dicer, a key enzyme in microRNA biogenesis (reviewed in (Sotillo and Thomas-Tikhonenko, 2011)). Beyond the hnRNPA1-Dicer axis, we identified strong (ΔPSI>50%) LSVs in ~30% of genes included in the COSMIC database because of their documented involvement in hematologic malignancies (Forbes et al., 2015; Futreal et al., 2004). Interestingly, four of these LSVs affect Drosha, another key enzyme in the microRNA biogenesis pathway (Sotillo and Thomas-Tikhonenko, 2011), suggesting a strong link between splicing and miRNA machineries.

Of even greater importance is the fact that B-ALL-specific LSVs affect 15 out of 20 top leukemia driver genes (including the aforementioned *FLT3*, *IL7R*, and *TP53*), with frequencies far exceeding those of somatic mutations. This discovery could explain why the prevalence of somatic mutations and copy number variations in B-ALL is low compared to other human cancers. It remains to be determined whether splicing alterations in oncogenes and tumor suppressor genes are functionally equivalent to gain-of-function and loss-of-function mutations, respectively. If so, interfering with splicing using RNA-based therapeutics and/or available small molecule inhibitors could be used to inhibit oncogenes such as *FLT3* and to activate dormant tumor suppressor gene such as *TP53*. Such strategies could yield tangible therapeutic benefits across a broad spectrum of childhood B-ALL subtypes.

## ACKNOWLEDGMENTS

This work was supported by grants from the NIH (T32 HL007439 to KLB, T32 CA 115299 to EG, K08 CA184418 to SKT, R01 AG046544 to YB), Stand Up To Cancer - St. Baldrick’s Pediatric Dream Team Translational Research Grant (SU2C-AACR-DT1113 to ATT), William Lawrence and Blanche Hughes Foundation (2016 Research Grant ATT), St. Baldrick’s Foundation (RG 527717 to ATT), Alex’s Lemonade Stand Foundation (Innovation Award to ATT), and CURE Childhood Cancer (Basic Research Award to KLB).

B-ALL samples used in this submission were passed to investigators with a coded identifier and no protected health information (PHI). Basic demographic, treatment, relapse, and survival outcome data for delivered specimens were provided through an online honest broker system. Specimens were obtained via informed consent on institutional research protocols in accordance with the Declaration of Helsinki.

The original RNA-Seq data sets will be deposited in the NCBI GEO database (accession number pending).

## Supplementary Materials for

Aberrant splicing in B-cell acute lymphoblastic leukemia by Black, A.S.Naqvi, et al.

**Supp. Fig. 1.**
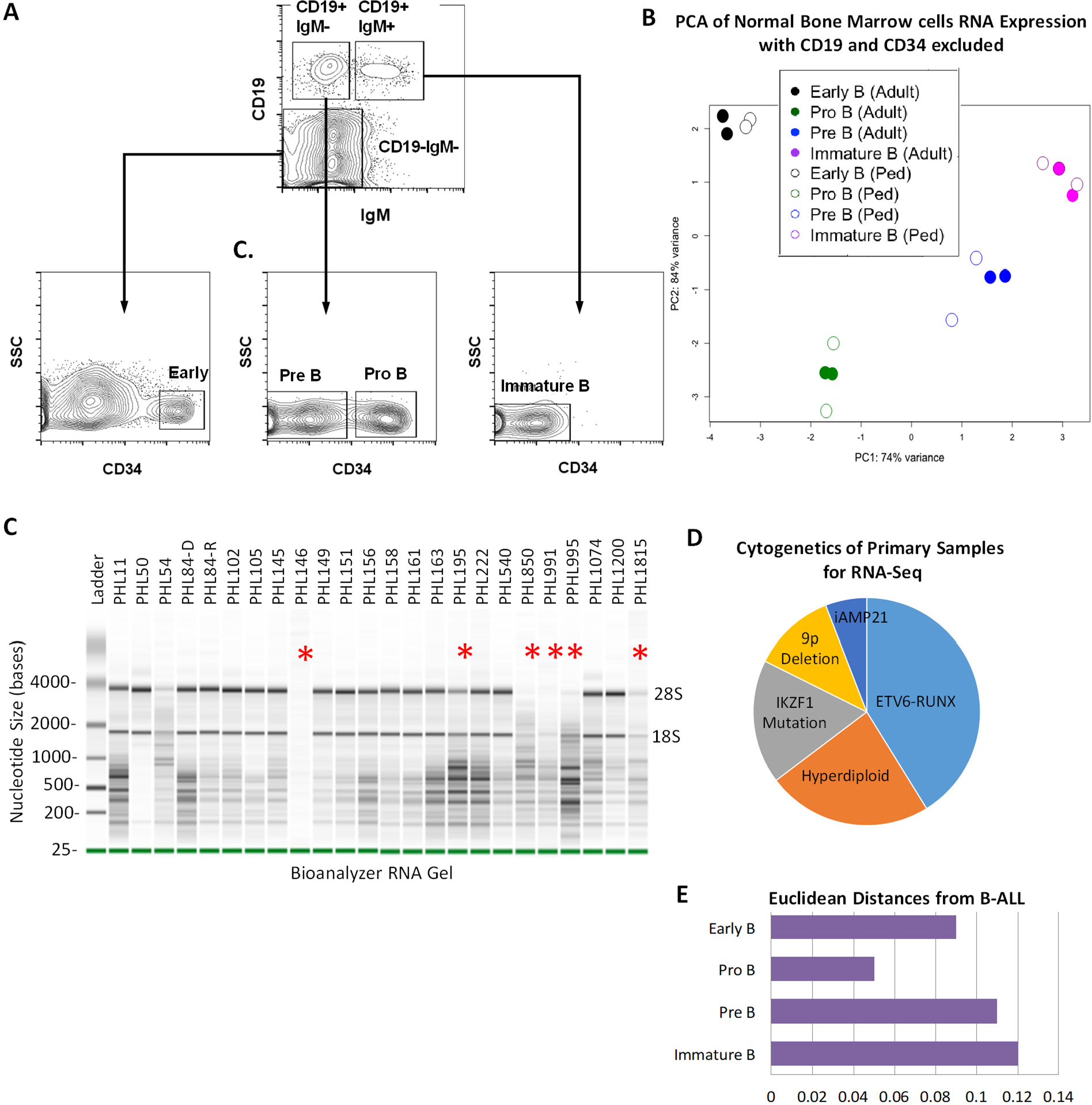
a. Lymphocytes were isolated from normal whole bone marrow aspirates and fractionated by flow cytometry into CD34+/CD19-/IgM-, CD34+/CD19+/IgM-, CD34-/CD19+/ IgM-, and CD34-/CD19+/IgM+ populations. b. PCA on RNA expression from 4 fractionated bone marrow samples without consideration of the expression of CD19 and CD34. c. RNA integrity (RIN) analysis of 24 cryopreserved leukemia samples. 18 of them passing QC criteria were chosen for Next Generation sequencing. Discarded samples are denoted with red asterisks. d. Pie chart showing distribution of cytogenetic abnormalities in 18 B-ALL samples. e. Euclidean distances showing closest clustering of B-ALL with pro-B cell fractions from PCA shown in Fig 1d.

**Supp. Fig. 2.**
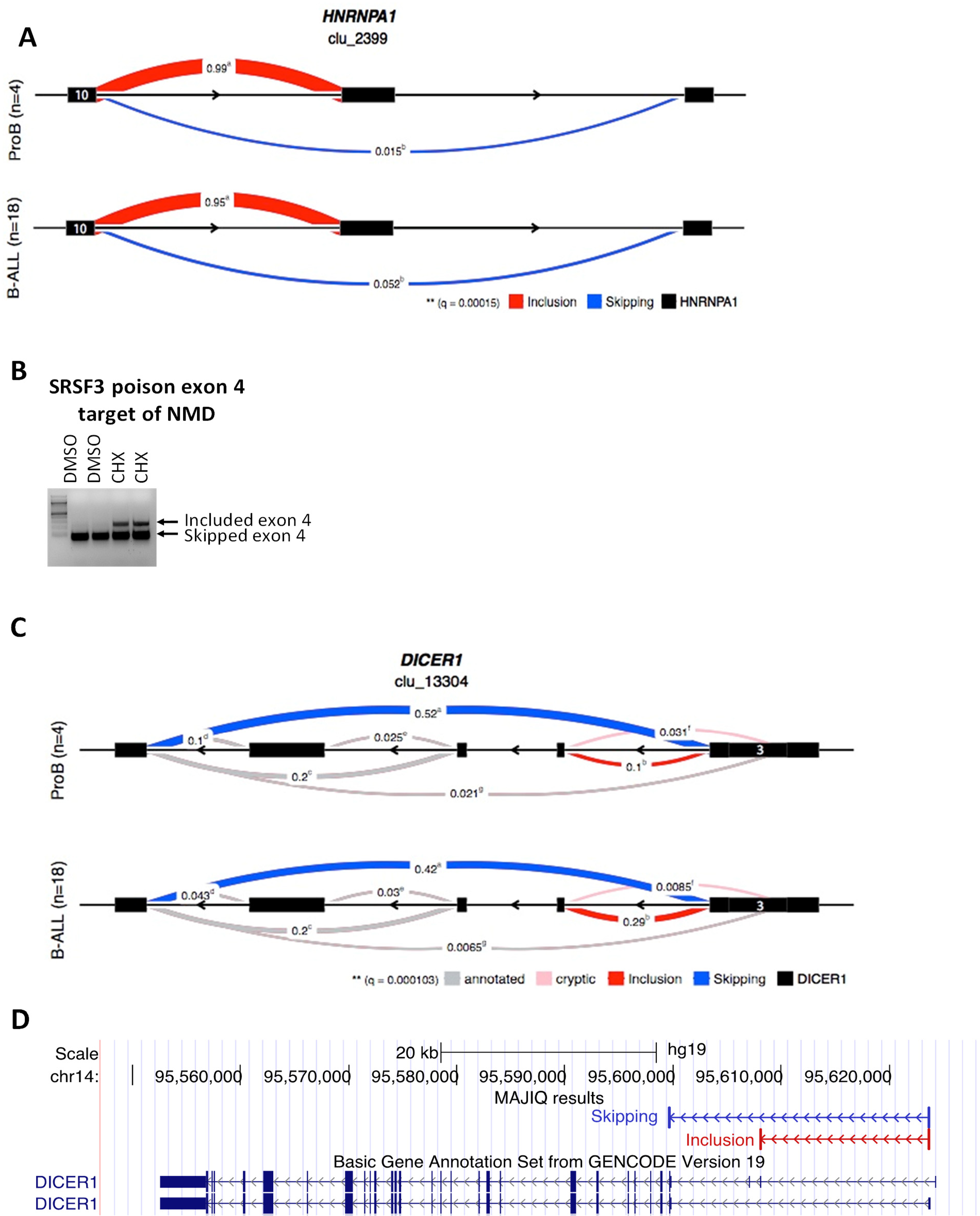
a. Detection of alternative splicing of hnRNPA1 3’ UTR by LeafCutter. LeafCutter visualization was adapted to highlight overlap with MAJIQ results. Red trace indicates inclusion of long, proximal exon 11. Blue trace indicates skipping to short, distal exon 12. b. PCR with primers detecting skipping or inclusion of SRSF3 poison exon 4 as a control for inhibition of NMD with cyclohexamide. c. Detection of alternative splicing of DICER1 by LeafCutter. d. UCSC Genome Browser track of DICER1 with MAJIQ LSVs indicated above.

**Supp. Fig. 3.**
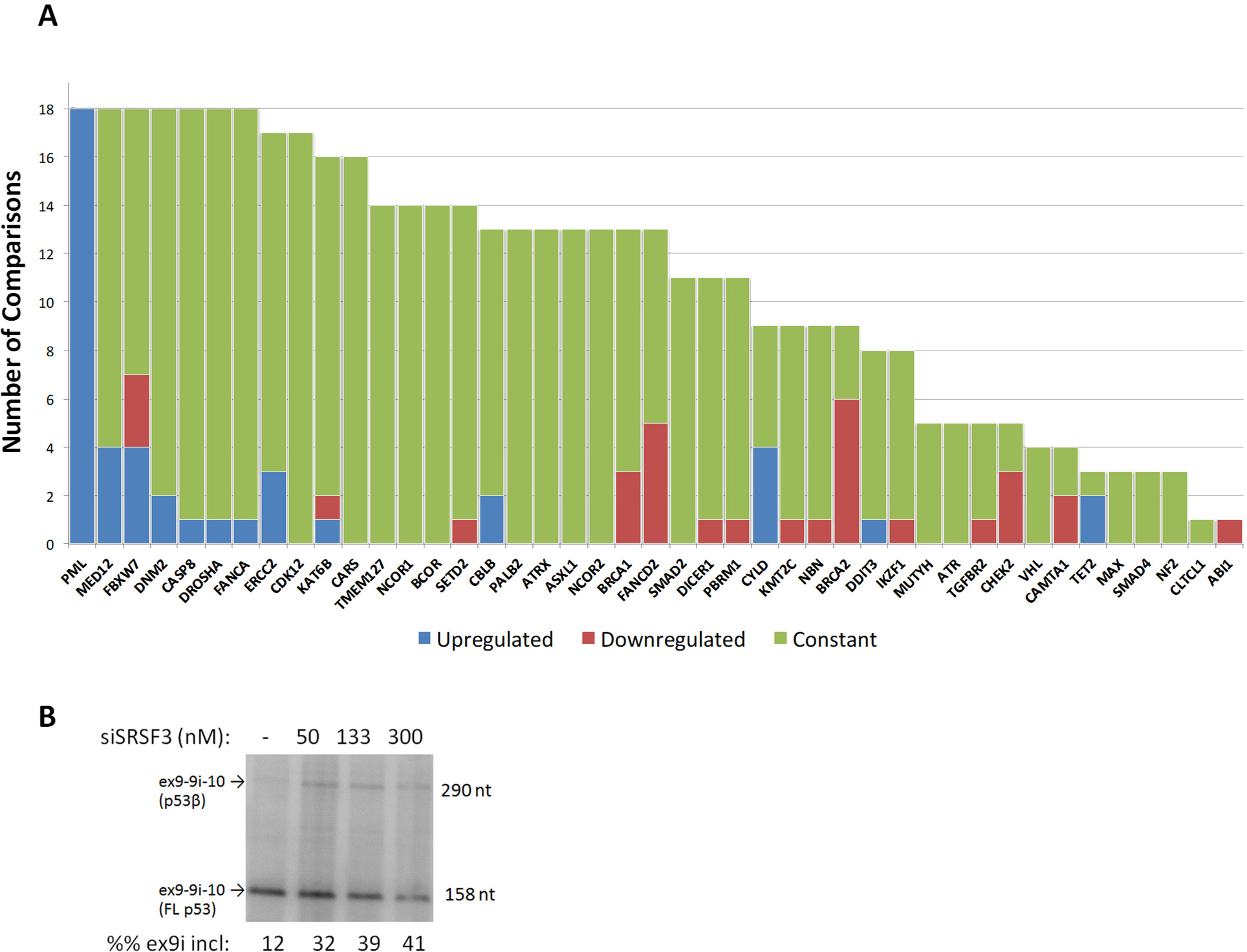
a. Differential expression analysis of COSMIC genes containing LSVs detected in primary B-ALL samples. Majority of samples showed constant expression of indicated genes compared to normal pro-B samples (green). Several samples showed downregulation (red) or upregulation (blue) of a few genes. b. Validation of alternative splicing of TP_53_ exon 9i in Nalm6 cells by radioactive PCR following siRNA mediated knockdown of SRSF3 at indicated concentrations.

## REFERENCES

Alderton, G.K. 2015. Immunotherapy: Skipping out epitopes. Nat Rev Cancer 15:699–699.

Amrani, N., R. Ganesan, S. Kervestin, D.A. Mangus, S. Ghosh, and A. Jacobson. 2004. A faux 3’-UTR promotes aberrant termination and triggers nonsense-mediated mRNA decay. Nature 432:112–118.

Behjati, S. 2015. Hiding from the enemy. Science Transl Med 7:313ec193–313ec193.

Blake, J.A., M. Dolan, H. Drabkin, D.P. Hill, N. Li, D. Sitnikov, S. Bridges, S. Burgess, T. Buza, F. McCarthy, et al. 2013. Gene Ontology annotations and resources. Nucleic Acids Res 41:D530–535.

Bourdon, J.C., K. Fernandes, F. Murray-Zmijewski, G. Liu, A. Diot, D.P. Xirodimas, M.K. Saville, and D.P. Lane. 2005. p53 isoforms can regulate p53 transcriptional activity. Genes Dev 19:2122–2137.

Burd, C.G., and G. Dreyfuss. 1994. RNA binding specificity of hnRNP A1: significance of hnRNP A1 high-affinity binding sites in pre-mRNA splicing. EMBO J 13:1197–1204.

Carter, M.S., S. Li, and M.F. Wilkinson. 1996. A splicing-dependent regulatory mechanism that detects translation signals. EMBO J 15:5965–5975.

Casero, D., S. Sandoval, C.S. Seet, J. Scholes, Y. Zhu, V.L. Ha, A. Luong, C. Parekh, and G.M. Crooks. 2015. Long non-coding RNA profiling of human lymphoid progenitor cells reveals transcriptional divergence of B cell and T cell lineages. Nat Immunol 16:1282–1291.

David, C.J., M. Chen, M. Assanah, P. Canoll, and J.L. Manley. 2010. HnRNP proteins controlled by c-Myc deregulate pyruvate kinase mRNA splicing in cancer. Nature 463:364–368.

Dennis, G., Jr., B.T. Sherman, D.A. Hosack, J. Yang, W. Gao, H.C. Lane, and R.A. Lempicki. 2003. DAVID: Database for Annotation, Visualization, and Integrated Discovery. Genome Biol 4:3.

Dobin, A., C.A. Davis, F. Schlesinger, J. Drenkow, C. Zaleski, S. Jha, P. Batut, M. Chaisson, and T.R. Gingeras. 2013. STAR: ultrafast universal RNA-seq aligner. Bioinformatics 29:15–21.

Dvinge, H., R.E. Ries, J.O. Ilagan, D.L. Stirewalt, S. Meshinchi, and R.K. Bradley. 2014. Sample processing obscures cancer-specific alterations in leukemic transcriptomes. Proc Natl Acad Sci USA 111:16802–16807.

Forbes, S.A., D. Beare, P. Gunasekaran, K. Leung, N. Bindal, H. Boutselakis, M. Ding, S. Bamford, C. Cole, S. Ward, et al. 2015. COSMIC: exploring the world’s knowledge of somatic mutations in human cancer. Nucleic Acids Res 43:D805–811.

Foulkes, W.D., J.R. Priest, and T.F. Duchaine. 2014. DICER1: mutations, microRNAs and mechanisms. Nat Rev Cancer 14:662–672.

Fry, T.J., N.N. Shah, R.J. Orentas, M. Stetler-Stevenson, C.M. Yuan, S. Ramakrishna, P. Wolters, S. Martin, C. Delbrook, B. Yates, et al. 2017. CD22-targeted CAR T cells induce remission in B-ALL that is naive or resistant to CD19-targeted CAR immunotherapy. Nature Med 10.1038/nm.4441.

Fujita, K., A.M. Mondal, I. Horikawa, G.H. Nguyen, K. Kumamoto, J.J. Sohn, E.D. Bowman, E.A. Mathe, A.J. Schetter, S.R. Pine, et al. 2009. p53 isoforms Δ133p53 and p53β are endogenous regulators of replicative cellular senescence. Nat Cell Biol 11:1135–1142.

Futreal, P.A., L. Coin, M. Marshall, T. Down, T. Hubbard, R. Wooster, N. Rahman, and M.R. Stratton. 2004. A census of human cancer genes. Nat Rev Cancer 4:177–183.

Gardner, R., D. Wu, S. Cherian, M. Fang, L.A. Hanafi, O. Finney, H. Smithers, M.C. Jensen, S.R. Riddell, D.G. Maloney, et al. 2016. Acquisition of a CD19-negative myeloid phenotype allows immune escape of MLL-rearranged B-ALL from CD19 CAR-T-cell therapy. Blood 127:2406–2410.

Graubert, T.A., D. Shen, L. Ding, T. Okeyo-Owuor, C.L. Lunn, J. Shao, K. Krysiak, C.C. Harris, D.C. Koboldt, D.E. Larson, et al. 2012. Recurrent mutations in the U2AF1 splicing factor in myelodysplastic syndromes. Nat Genet 44:53–U77.

Gu, Z., M. Churchman, K. Roberts, Y. Li, Y. Liu, R.C. Harvey, K. McCastlain, S.C. Reshmi, D. Payne-Turner, I. Iacobucci, et al. 2016. Genomic analyses identify recurrent MEF2D fusions in acute lymphoblastic leukaemia. Nat Commun 7:13331.

Haso, W., D.W. Lee, N.N. Shah, M. Stetler-Stevenson, C.M. Yuan, I.H. Pastan, D.S. Dimitrov, R.A. Morgan, D.J. FitzGerald, D.M. Barrett, et al. 2013. Anti-CD22-chimeric antigen receptors targeting B-cell precursor acute lymphoblastic leukemia. Blood 121:1165–1174.

Huang, L., C.H. Lou, W. Chan, E.Y. Shum, A. Shao, E. Stone, R. Karam, H.W. Song, and M.F. Wilkinson. 2011. RNA homeostasis governed by cell type-specific and branched feedback loops acting on NMD. Mol Cell 43:950–961.

Hunger, S.P., M.L. Loh, J.A. Whitlock, N.J. Winick, W.L. Carroll, M. Devidas, E.A. Raetz, and C.O.G.A.L.L. Committee. 2013. Children’s Oncology Group’s 2013 blueprint for research: acute lymphoblastic leukemia. Pediatr Blood Cancer 60:957–963.

Jacoby, E., S.M. Nguyen, T.J. Fountaine, K. Welp, B. Gryder, H. Qin, Y. Yang, C.D. Chien, A.E. Seif, H. Lei, et al. 2016. CD19 CAR immune pressure induces B-precursor acute lymphoblastic leukaemia lineage switch exposing inherent leukaemic plasticity. Nat Commun 7:12320.

Jumaa, H., and P.J. Nielsen. 1997. The splicing factor SRp20 modifies splicing of its own mRNA and ASF/SF2 antagonizes this regulation. EMBO J 16:5077–5085.

Li, Y.I., D.A. Knowles, J. Humphrey, A.N. Barbeira, S.P. Dickinson, H.K. Im, and J.K. Pritchard. 2018. Annotation-free quantification of RNA splicing using LeafCutter. Nat Genet 50:151–158.

Love, M.I., W. Huber, and S. Anders. 2014. Moderated estimation of fold change and dispersion for RNA-seq data with DESeq2. Genome Biol 15:550.

Maino, E., M. Bonifacio, A.M. Scattolin, and R. Bassan. 2016. Immunotherapy approaches to treat adult acute lymphoblastic leukemia. Expert Rev Hematol 9:563–577.

Marcel, V., K. Fernandes, O. Terrier, D.P. Lane, and J.C. Bourdon. 2014. Modulation of p53? and p53? expression by regulating the alternative splicing of TP53 gene modifies cellular response. Cell Death Differ 21:1377–1387.

Matera, A.G., and Z. Wang. 2014. A day in the life of the spliceosome. Nat Rev Mol Cell Biol 15:108–121.

Maude, S.L., N. Frey, P.A. Shaw, R. Aplenc, D.M. Barrett, N.J. Bunin, A. Chew, V.E. Gonzalez, Z. Zheng, S.F. Lacey, et al. 2014. Chimeric antigen receptor T cells for sustained remissions in leukemia. N Engl J Med 371:1507–1517.

Michlewski, G., and J.F. Caceres. 2010. Antagonistic role of hnRNP A1 and KSRP in the regulation of let-7a biogenesis. Nat Struct Mol Biol 17:1011–1018.

Mullighan, C.G., S. Goorha, I. Radtke, C.B. Miller, E. Coustan-Smith, J.D. Dalton, K. Girtman, S. Mathew, J. Ma, S.B. Pounds, et al. 2007. Genome-wide analysis of genetic alterations in acute lymphoblastic leukaemia. Nature 446:758–764.

Norton, S., J. Vaquero-Garcia, N.F. Lahens, G.R. Grant, and Y. Barash. 2017. Outlier detection for improved differential splicing quantification from RNA-Seq experiments with replicates. Bioinformatics

Pajic, A., M.S. Staege, D. Dudziak, M. Schuhmacher, D. Spitkovsky, G. Eissner, M. Brielmeier, A. Polack, and G.W. Bornkamm. 2001. Antagonistic effects of c-myc and Epstein-Barr virus latent genes on the phenotype of human B cells. Int J Cancer 93:810–816.

Papaemmanuil, E., M. Cazzola, J. Boultwood, L. Malcovati, P. Vyas, D. Bowen, A. Pellagatti, J.S. Wainscoat, E. Hellstrom-Lindberg, C. Gambacorti-Passerini, et al. 2011. Somatic SF3B1 Mutation in Myelodysplasia with Ring Sideroblasts. N Engl J Med 365:1384–1395.

Paz, I., I. Kosti, M. Ares, Jr., M. Cline, and Y. Mandel-Gutfreund. 2014. RBPmap: a web server for mapping binding sites of RNA-binding proteins. Nucleic Acids Res 42:W361–367.

Quesada, V., L. Conde, N. Villamor, G.R. Ordonez, P. Jares, L. Bassaganyas, A.J. Ramsay, S. Bea, M. Pinyol, A. Martinez-Trillos, et al. 2012. Exome sequencing identifies recurrent mutations of the splicing factor SF3B1 gene in chronic lymphocytic leukemia. Nat Genet 44:47–52.

Raetz, E.A., M.S. Cairo, M.J. Borowitz, S.M. Blaney, M.D. Krailo, T.A. Leil, J.M. Reid, D.M. Goldenberg, W.A. Wegener, W.L. Carroll, et al. 2008. Chemoimmunotherapy reinduction with epratuzumab in children with acute lymphoblastic leukemia in marrow relapse: a Children’s Oncology Group Pilot Study. J Clin Oncol 26:3756–3762.

Ray, D., H. Kazan, K.B. Cook, M.T. Weirauch, H.S. Najafabadi, X. Li, S. Gueroussov, M. Albu, H. Zheng, A. Yang, et al. 2013. A compendium of RNA-binding motifs for decoding gene regulation. Nature 499:172–177.

Roberts, K.G., and C.G. Mullighan. 2015. Genomics in acute lymphoblastic leukaemia: insights and treatment implications. Nat Rev Clin Oncol 12:344–357.

Rodriguez-Aguayo, C., P.D.C. Monroig, R.S. Redis, E. Bayraktar, M.I. Almeida, C. Ivan, E. Fuentes-Mattei, M.H. Rashed, A. Chavez-Reyes, B. Ozpolat, et al. 2017. Regulation of hnRNPA1 by microRNAs controls the miR-18a-K-RAS axis in chemotherapy-resistant ovarian cancer. Cell Discov 3:17029.

Scheuermann, R.H., and E. Racila. 1995. CD19 antigen in leukemia and lymphoma diagnosis and immunotherapy. Leuk Lymphoma 18:385–397.

Sebestyen, E., B. Singh, B. Minana, A. Pages, F. Mateo, M.A. Pujana, J. Valcarcel, and E. Eyras. 2016. Large-scale analysis of genome and transcriptome alterations in multiple tumors unveils novel cancer-relevant splicing networks. Genome Res 26:732–744.

Sikaria, S., I. Aldoss, and M. Akhtari. 2016. Monoclonal antibodies and immune therapies for adult precursor B-acute lymphoblastic leukemia. Immunol Lett 172:113–123.

Sotillo, E., D.M. Barrett, K.L. Black, A. Bagashev, D. Oldridge, G. Wu, R. Sussman, C. Lanauze, M. Ruella, M.R. Gazzara, et al. 2015. Convergence of acquired mutations and alternative splicing of CD19 enables resistance to CART-19 immunotherapy. Cancer Discov 5:1282–1295.

Sotillo, E., and A. Thomas-Tikhonenko. 2011. Shielding the messenger (RNA): microRNA-based anticancer therapies. Pharmacol Ther 131:18–32.

Tang, Y., I. Horikawa, M. Ajiro, A.I. Robles, K. Fujita, A.M. Mondal, J.K. Stauffer, Z.M. Zheng, and C.C. Harris. 2013. Downregulation of splicing factor SRSF3 induces p53beta, an alternatively spliced isoform of p53 that promotes cellular senescence. Oncogene 32:2792–2798.

Tasian, S.K., M.Y. Doral, M.J. Borowitz, B.L. Wood, I.M. Chen, R.C. Harvey, J.M. Gastier-Foster, C.L. Willman, S.P. Hunger, C.G. Mullighan, et al. 2012. Aberrant STAT5 and PI3K/mTOR pathway signaling occurs in human CRLF2-rearranged B-precursor acute lymphoblastic leukemia. Blood 120:833–842.

Topp, M.S., N. Gokbuget, A.S. Stein, G. Zugmaier, S. O’Brien, R.C. Bargou, H. Dombret, A.K. Fielding, L. Heffner, R.A. Larson, et al. 2015. Safety and activity of blinatumomab for adult patients with relapsed or refractory B-precursor acute lymphoblastic leukaemia: a multicentre, single-arm, phase 2 study. Lancet Oncol 16:57–66.

Vaquero-Garcia, J., A. Barrera, M.R. Gazzara, J. Gonzalez-Vallinas, N.F. Lahens, J.B. Hogenesch, K.W. Lynch, and Y. Barash. 2016. A new view of transcriptome complexity and regulation through the lens of local splicing variations. eLife 5:e11752.

Wang, L.L., M.S. Lawrence, Y.Z. Wan, P. Stojanov, C. Sougnez, K. Stevenson, L. Werner, A. Sivachenko, D.S. DeLuca, L. Zhang, et al. 2011. SF3B1 and Other Novel Cancer Genes in Chronic Lymphocytic Leukemia. N Engl J Med 365:2497–2506.

Weighardt, F., G. Biamonti, and S. Riva. 1996. The roles of heterogeneous nuclear ribonucleoproteins (hnRNP) in RNA metabolism. Bioessays 18:747–756.

Yoshida, K., M. Sanada, Y. Shiraishi, D. Nowak, Y. Nagata, R. Yamamoto, Y. Sato, A. Sato-Otsubo, A. Kon, M. Nagasaki, et al. 2011. Frequent pathway mutations of splicing machinery in myelodysplasia. Nature 478:64–69.

Zerbino, D.R., P. Achuthan, W. Akanni, M.R. Amode, D. Barrell, J. Bhai, K. Billis, C. Cummins, A. Gall, C.G. Giron, et al. 2017. Ensembl 2018. Nucleic Acids Res

